# *In Silico* analysis of the structural and functional impact of deleterious nsSNPs in the human *RETREG1* gene associated with congenital sensory neuropathy type II

**DOI:** 10.64898/2026.01.02.697353

**Authors:** Mohammed Ezzeldein Mohammed Alsied, Tebyan Ameer Abdelhameed Abbas, Seedahmed A. Mohammed

## Abstract

**Background:** Mutations in the *RETREG1* gene are known to cause Hereditary Sensory and Autonomic Neuropathy type II (HSAN II), a severe congenital disorder affecting sensory neurons. However, the full spectrum of pathogenic single nucleotide polymorphisms (SNPs) and their specific structural consequences remain incompletely characterized.

**Objectives:** This study aimed to elucidate pathogenic nsSNPs and their role in the Congenital Sensory Neuropathy (HSAN II) by employing *in-silico* analysis.

**Method:** The nsSNPs of *RETREG1* were retrieved from the dbSNP in NCBI database. Different *in silico* tools, SIFT, PolyPhen-2, SNP&GO, PHD-SNP, SNAP2, I-mutant, Project Hope, MutPred, ConSurf, phyre2, Chimera, and GeneMANIA, were used for predicting the pathogenicity, protein stability, evolutionary conservation, structural alterations, and protein–protein interaction networks.for RETREG1 gene.

**Result:** Five nsSNPs were identified as “damaging” or deleterious by using the above software (Y221C, G216R, G211R, L119V, W107C). Four SNPs (Y221C, G211R, L119V, and W107C) were predicted to decrease protein stability, while the fifth SNP (G216R) was expected to increase it. Structural modeling revealed that all five mutations are located within the critical Reticulon Homology Domain (RHD), where they are predicted to cause steric clashes, disrupt hydrophobic packing, and impair protein-membrane interactions.

**Conclusion:** This integrated *in silico* analysis identifies four novel deleterious nsSNPs in RETREG1 (W107C, L119V, G211R, Y221C) and confirms the established G216R variant. These mutations are predicted to impair RETREG1 structure and function, providing mechanistic insight into HSAN II pathogenesis and prioritizing candidates for future experimental validation.

## Introduction

Hereditary sensory autonomic neuropathy (HSAN) comprise a clinically and genetically heterogeneous group of disorders primarily affecting sensory and autonomic neurons (Rudnik-Schöneborn et al., 2017; Eggermann et al., 2018; Low et al., 2003; Verpoorten et al., 2006). It includes five different types: HSAN I, II, III, IV, and V (Axelrod & Gold-von Simson, 2007; Ota et al., 1973).

HSAN type II is an autosomal recessive form characterized by early onset and gradual distal sensory loss across all modalities frequently, frequently resulting in severe consequences such as recurrent ulcerations, soft tissue infections, and osteomyelitis, which may necessitate amputation of the distal extremities (Axelrod & Gold-von Simson, 2007; Rudnik-Schöneborn et al., 2017). HSAN type II is genetically diverse and can be further subdivided based on the causal gene, includeing *KIF1A*, *SCN9A, WNK1 and RETREG1* (formerly known as *FAM134B*) (Falcão de Campos et al., 2019).

Mutations in the RETREG1 gene are specifically associated with the HSAN IIB subtype (Islam et al., 2018). The RETREG protein functions as a Cis-Golgi transmembrane endoplasmic reticulum (ER) receptor and is a key regulator of ER-phagy through its reticulon-homology domain (RHD) and LC3-interacting region (LIR) (Popelka & Klionsky, 2020).

While loss-of-function mutations in RETREG1 have been linked to the disease, the precise molecular mechanisms by which specific genetic variants, particularly missense mutations, disrupt protein function and lead to neuropathy remain largely uncharacterized (Islam et al., 2018; Wakil et al., 2018).

Single nucleotide polymorphisms (SNPs) are the most common form of genetic variation. Non-synonymous SNPs (nsSNPs) that result in amino acid substitutions can be particularly deleterious by altering protein stability, interaction interfaces, and function. nsSNP structural and functional analyses may aid in the creation of personalized medicine based on genetic variation (Hussain et al., 2012).

In this study, we performed a comprehensive *in silico* analysis to predict the structural and functional consequences of nsSNPs in the human RETREG1 gene. Using a combination of sequence-based, evolutionary, and structure-based prediction tools, we identified high-risk deleterious variants, assessed their impact on protein stability, and modeled the structural changes induced by these mutations. Our findings aim to provide insights into the molecular pathology of HSAN IIB and identify key residues critical for RETREG1 function

## Materials and Methods

### Retrieval of protein and SNP data

The National Center for Biotechnology Information (NCBI) website was used to retrieve data of the RETREG1 gene. Information on SNPs (single nucleotide polymorphisms) was obtained from the NCBI dbSNP database (https://www.ncbi.nlm.nih.gov/projects/SNP/). A total of 345 missense nsSNPs were selected for further computational analysis. From UniProt (https://www.uniprot.org), the gene ID and sequence were retrieved (Uniprot ID: Q9H6L5).

### Prediction of pathogenic nsSNPs

To predict the deleterious effects of the identified nsSNPs, we employed a suite of computational tools based on different algorithms.

### SIFT

SIFT (Sorting Intolerant From Tolerant) was used to distinguish between intolerant and tolerant amino acid substitutions, and to determine whether a protein is likely to exhibit phenotypic changes as a result of an amino acid substitution by considering the position of the mutation and the type of amino acid change. The substitution is expected to be deleterious if its standardized value falls below a threshold. according to SIFT scores, Intolerant or deleterious amino acid substitutions are expected to have SIFT scores <0.05, whereas tolerant amino acid substitutions have values >0.05 ((Kumar et al., 2009; Ng et al., 2003). The tool is available at (http://sift.jcvi.org/www/SIFT_seq_submit2.html).

### PolyPhen-2

Accessible at (http://genetics.bwh.harvard.edu/pph2), Polymorphism Phenotyping v2 was used to predict the impact of amino acid substitutions on the structure and function of the protein. Position-specific independent count (PSIC) scores were calculated for each variant, along with the difference between the two variants’ PSIC scores. The functional effect was considered greater when the difference in PSIC scores was large. According to PSIC (PolyPhen) scores there are three categories for each variant: probably damaging (0.85-1), possibly damaging (0.2-0.85), and benign (0.00–0.2) (Adzhubei et al., 2013)

### SNAP2

SNAP2 was used to distinguish between effect and neutral variants/non-synonymous SNPs using neural networks. The tool is accessible at http://www.rostlab.org/services/SNAP (Bromberg & Rost, 2007)

### PHD-SNP

PHD-SNP was used to predict the associated variation as either be disease-related (Disease) or neutral polymorphic (Neutral) using Support Vector Machines (SVM)-based approach. The tool is accessible at http://snps.biofold.org/phd-snp/phd-snp.html (Capriotti et al., 2006)

### SNP & GO

SNP & GO was used for predicting single point protein mutations that are most likely to be responsible for human disease (Capriotti et al., 2017). Available at (http://snps.biofold.org/snps-and-go/snps-and-go.html).

A nsSNP was considered high-risk and selected for further analysis if it was predicted to be deleterious by the majority of these tools.

### Analysis of Protein Stability

#### I–Mutant

Accessible at (https://folding.biofold.org/i-mutant/i-mutant2.0.html). It was used to predict how a protein’s stability would change in response to high-risk nsSNPs. This tool predicts the change in Gibbs free energy (ΔΔG) upon mutation; a negative ΔΔG value indicates a decrease in protein stability (Essa et al., 2019)

### Evolutionary Conservation Analysis

The evolutionary conservation of amino acid residues was analyzed using the ConSurf server (https://consurf.tau.ac.il). This server calculates conservation scores based on the phylogenetic relationships between homologous sequences, identifying functionally important residues

### Protein Structure Modeling and Analysis

The three-dimensional structure of the wild-type RETREG1 protein was modeled using the PHYRE2 server (Protein Homology/analogY Recognition Engine V 2.0; http://www.sbg.bio.ic.ac.uk/phyre2) in intensive mode (Goswami, 2015). The high-risk mutant structures were generated by introducing the respective amino acid changes into the wild-type model using UCSF Chimera (v1.16; https://www.cgl.ucsf.edu/chimera) (Mustafa et al., 2020). Structural comparisons between wild-type and mutant models were performed to visualize changes. Further structural analysis, including the visualization of domain architecture and mutant effects, was conducted using Project HOPE (https://www3.cmbi.umcn.nl/hope).

### Prediction of Functional Implications

The molecular mechanistic impact of the high-risk mutations was predicted using MutPred2 (http://mutpred.mutdb.org/), which provides hypotheses on the gain or loss of structural and functional properties due to the amino acid substitution (Pejaver et al., 2020)

### Protein Interaction Network Analysis

A protein-protein interaction network for RETREG1 was generated using **GeneMANIA** (http://www.genemania.org) to identify functional partners and potential pathways affected by the mutations(Abbas et al., 2019)

## Results

### Identification and Prioritization of High-Risk nsSNPs in RETREG1

It’s important to understand the consequences of harmful nsSNPs in RETREG1 and how these relate to various disorders and diseases. Here, we used computational analysis to determine which nsSNPs were the most deleterious and how they may affect the structure and function of the RETREG1 protein. A total of 345 nsSNPs were retrieved from dbSNP in NCBI database. To identify potentially deleterious variants, we employed a multi-step computational filtering approach using three distinct prediction tools: SIFT, PolyPhen-2, and SNAP2.

Of the 345 variants, 68 were predicted as a DAMAGE substitution by SIFT, 56 were predicted as “effect” by SNAP2, and by using PolyPhen2, 10 SNPs predicted as possibly damaging, and 28 as a probably damaging (Table, 1).

**Table 1:** Prediction of functional effect of deleterious and damaging nsSNPs of RETREG1 by SIFT, polyphene2, and SNAP2.

| Mutation | SIFT pre | SIFT score | PPH2 pre | PPH2 score | SNAP2 pre | SNAP2 score |
| --- | --- | --- | --- | --- | --- | --- |
| G496D | DAMAGE | 0.00 | PROBABLY DAMAGING | 0.990 | Effect | 20 |
| N492D | DAMAGE | 0.00 | POSSIBLY DAMAGING | 0.651 | Effect | 17 |
| S491L | DAMAGE | 0.00 | PROBABLY DAMAGING | 0.990 | Effect | 55 |
| G488S | DAMAGE | 0.00 | PROBABLY DAMAGING | 1.000 | Effect | 4 |
| D475H | DAMAGE | 0.00 | POSSIBLY DAMAGING | 0.855 | Effect | 10 |
| G471A | DAMAGE | 0.00 | PROBABLY DAMAGING | 0.991 | Effect | 29 |
| L458R | DAMAGE | 0.00 | PROBABLY DAMAGING | 0.999 | Effect | 72 |
| L457P | DAMAGE | 0.00 | PROBABLY DAMAGING | 1.000 | Effect | 68 |
| E456D | DAMAGE | 0.00 | PROBABLY DAMAGING | 0.997 | Effect | 56 |
| D446Y | DAMAGE | 0.01 | PROBABLY DAMAGING | 0.999 | Effect | 20 |
| G358V | DAMAGE | 0.01 | PROBABLY<br>DAMAGING | 0.997 | Effect | 1 |
| N355Y | DAMAGE | 0.03 | PROBABLY<br>DAMAGING | 1.000 | Effect | 17 |
| S352F | DAMAGE | 0.01 | PROBABLY<br>DAMAGING | 0.994 | Effect | 28 |
| P351S | DAMAGE | 0.00 | PROBABLY<br>DAMAGING | 0.999 | Effect | 19 |
| D336E | DAMAGE | 0.00 | PROBABLY<br>DAMAGING | 0.999 | Effect | 14 |
| L335V | DAMAGE | 0.00 | POSSIBLY<br>DAMAGING | 0.824 | Effect | 30 |
| D334E | DAMAGE | 0.00 | PROBABLY<br>DAMAGING | 0.999 | Effect | 21 |
| T329I | DAMAGE | 0.00 | PROBABLY<br>DAMAGING | 0.999 | Effect | 9 |
| T326P | DAMAGE | 0.00 | PROBABLY<br>DAMAGING | 0.988 | Effect | 16 |
| S312F | DAMAGE | 0.00 | PROBABLY<br>DAMAGING | 0.999 | Effect | 26 |
| D284G | DAMAGE | 0.02 | PROBABLY<br>DAMAGING | 1.000 | Effect | 44 |
| I246S | DAMAGE | 0.01 | PROBABLY<br>DAMAGING | 0.970 | Effect | 64 |
| Y243H | DAMAGE | 0.00 | PROBABLY<br>DAMAGING | 0.995 | Effect | 73 |
| P231A | DAMAGE | 0.00 | PROBABLY<br>DAMAGING | 1.000 | Effect | 71 |
| Y221C | DAMAGE | 0.00 | PROBABLY<br>DAMAGING | 1.000 | Effect | 67 |
| G216R | DAMAGE | 0.00 | PROBABLY<br>DAMAGING | 1.000 | Effect | 90 |
| I214S | DAMAGE | 0.01 | PROBABLY<br>DAMAGING | 0.984 | Effect | 80 |
| G211R | DAMAGE | 0.00 | PROBABLY<br>DAMAGING | 1.000 | Effect | 94 |
| F196I | DAMAGE | 0.00 | PROBABLY<br>DAMAGING | 1.000 | Effect | 67 |
| P193S | DAMAGE | 0.00 | PROBABLY<br>DAMAGING | 1.000 | Effect | 71 |
| S192R | DAMAGE | 0.03 | POSSIBLY<br>DAMAGING | 0.801 | Effect | 32 |
| L119V | DAMAGE | 0.00 | PROBABLY<br>DAMAGING | 0.997 | Effect | 50 |
| W107C | DAMAGE | 0.00 | POSSIBLY<br>DAMAGING | 0.837 | Effect | 48 |
| N103S | DAMAGE | 0.00 | POSSIBLY<br>DAMAGING | 0.711 | Effect | 75 |
| S95R | DAMAGE | 0.000 | POSSIBLY<br>DAMAGING | 0.748 | Effect | 85 |
| P92L | DAMAGE | 0.00 | POSSIBLY<br>DAMAGING | 0.858 | Effect | 69 |
| R82H | DAMAGE | 0.04 | POSSIBLY<br>DAMAGING | 0.942 | Effect | 30 |
| W70R | DAMAGE | 0.00 | POSSIBLY<br>DAMAGING | 0.910 | Effect | 20 |

A consensus approach revealed 38 nsSNPs that were predicted to be deleterious by at least one tool. These 38 variants were selected for further analysis to identify those with the highest potential pathogenicity.

### Prediction of Disease-Association and Impact on Protein Stability

To further refine our list of high-risk candidates, the 38 deleterious nsSNPs were analyzed using two disease-specific predictors, PhD-SNP and SNP&GO. This analysis identified 5 nsSNPs (Y221C, G216R, G211R, L119V, and W107C) that were consistently predicted to be disease-associated by both tools (Table, 2), highlighting them as the most promising candidates linked to HSAN IIB pathology.

**Table 2:** Prediction of disease related and pathological effect of nsSNPs by PHD-SNP and SNP&GO.

| dbSNP rs# | Mutation | Prediction<br>SNP&GO | Probability<br>SNP&GO | RI | Prediction<br>PHD | Probability<br>PHD | RI |
| --- | --- | --- | --- | --- | --- | --- | --- |
| rs770744307 | Y221C | Disease | 0.58 | 1.00 | Disease | 0.88 | 8.00 |
| rs1579584286 | G216R | Disease | 0.65 | 3.00 | Disease | 0.92 | 8.00 |
| rs1738748948 | G211R | Disease | 0.68 | 4.00 | Disease | 0.91 | 8.00 |
| rs1467620688 | L119V | Disease | 0.51 | 0.00 | Disease | 0.75 | 5.00 |
| rs200065908 | W107C | Disease | 0.60 | 2.00 | Disease | 0.93 | 9.00 |

RI: Reliability Index. It is evaluated from the output of the support vector machine O. The threshold probability is 0.5. A disease associated nsSNPs have a probability scores above 0.5. The RI between 5 and 9 for PHD-SNP, and between 0 and 4 for SNP&GO are predicted as disease-associated nsSNPs as well.

### Prediction of change in RETREG1 stability due to mutation by I-mutant server

Protein structure and functional activity depends on protein stability (Shoichet et al., 1995). We next evaluated the impact of the five high-risk nSNPs on RETREG1 protein stability using I-Mutant. The majority of mutations (4 out of 5: Y221C, G211R, L119V, and W107C) were predicted to significantly decrease protein stability (ΔΔG < 0), suggesting a mechanism through which they could impair protein function. Interestingly, the G216R mutation was predicted to increase stability, which may also disrupt normal function by reducing conformational flexibility (Table, 3).

**Table 3:** prediction of nsSNPs effect on protein structure stability using I-mutant.

| dbSNP rs# | Mutation | I mutant probability | I mutant score |
| --- | --- | --- | --- |
| rs770744307 | Y221C | Decrease | -0.41 |
| rs1579584286 | G216R | Increase | 0.00 |
| rs1738748948 | G211R | Decrease | -0.67 |
| rs1467620688 | L119V | Decrease | -1.41 |
| rs200065908 | W107C | Decrease | -1.50 |

### Molecular Mechanistic and Evolutionary Conservation Analysis

The molecular consequences of the five high-risk mutations were predicted using MutPred2 (Table 4). The variants Y221C, G216R, and G211R—all located in a C-terminal region—were strongly predicted to cause gain of Helix and disrupt ordered protein interfaces.. The L119V was also predicted to cause altered ordered interface, while the W107C was predicted to affect a signal peptide, suggesting diverse mechanisms of dysfunction. Evolutionary conservation analysis using ConSurf revealed that the residues affected by four of the five mutations (G211, G216, Y221, and L119) are highly conserved (scores of 8-9 on a 1-9 scale), indicating strong negative selection pressure and implying their critical importance for RETREG1 function (Figure 1). The residue W107 showed moderate conservation (score of 4). These results show that some nsSNPs may account for potential structural and functional changes in RETREG1 protein (Table, 4)

**Figure 1:**
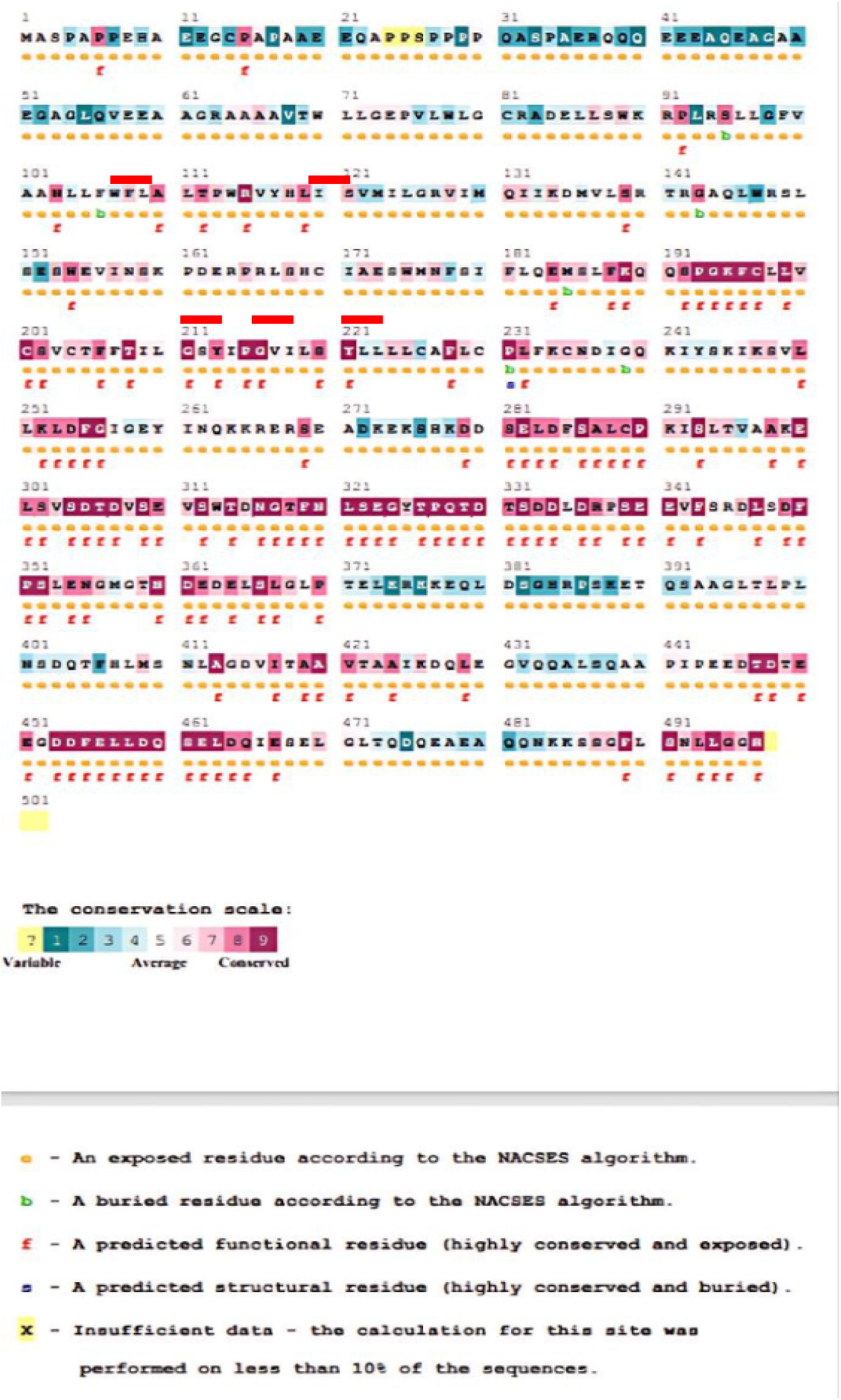
Evolutionary conservancy of RETREG1 produced by Consurf. Modelling of amino acid substitution on RETREG1 protein structure using chimera and phyre2. The red lines indicate the position of the novel nsSNPs

**Table 4:** Analysis of the effect of nsSNPs in RETREG1 structure, function, and evolution by MutPred server.

| dbSNP rs# | Mutation | MutPred probability | Feature |
| --- | --- | --- | --- |
| rs770744307 | Y221C | 0.843 | Altered Ordered interface (Pr = 0.36 \| P = 2.0e-03); Gain of Helix (Pr = 0.27 \| P = 0.04). |
| rs1579584286 | G216R | 0.834 | Gain of Helix (Pr = 0.34 \| P = 1.1e-03); Altered Ordered interface (Pr = 0.25 \| P = 0.02). |
| rs1738748948 | G211R | 0.883 | Gain of Helix (Pr = 0.33 \| P = 1.8e-03); Altered Ordered interface (Pr = 0.28 \| P = 5.4e-03). |
| rs1467620688 | L119V | 0.516 | Altered Ordered interface (Pr = 0.26 \| P = 0.01). |
| rs200065908 | W107C | 0.548 | Altered Signal peptide (Pr = 0.02 \| P = 4.8e-03). |

To elucidate the structural impact of the five high-risk nsSNPs, we generated a homology model of the RETREG1 RHD domain and modeled each mutation. All five variants (W107C, L119V, G211R, G216R, Y221C) are located within the core of the RHD, a domain essential for inducing ER membrane curvature and facilitating oligomerization.

Notably, the G211R and G216R mutations substitute a small, flexible glycine for a large, positively charged arginine. Structural analysis revealed that this substitution introduces significant steric clashes and is predicted to disrupt the precise packing of the transmembrane helices, likely hindering oligomerization (Figure 2 and 3).

**Figure 2:**
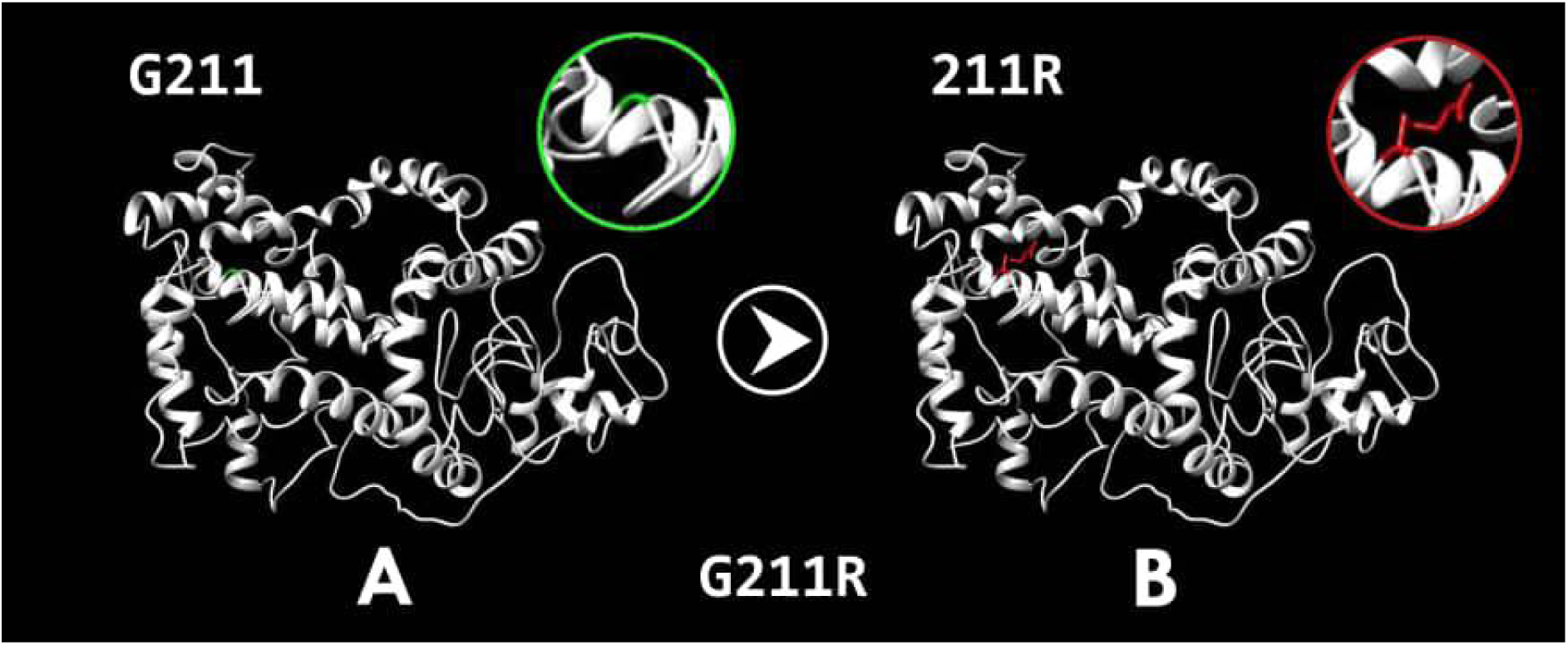
rs1738748948 (G211R). The amino acid Glycine in the wild protein (A) with green colour changed to Arginine in the mutant protein (B) with red colour at position 211.

**Figure 3:**
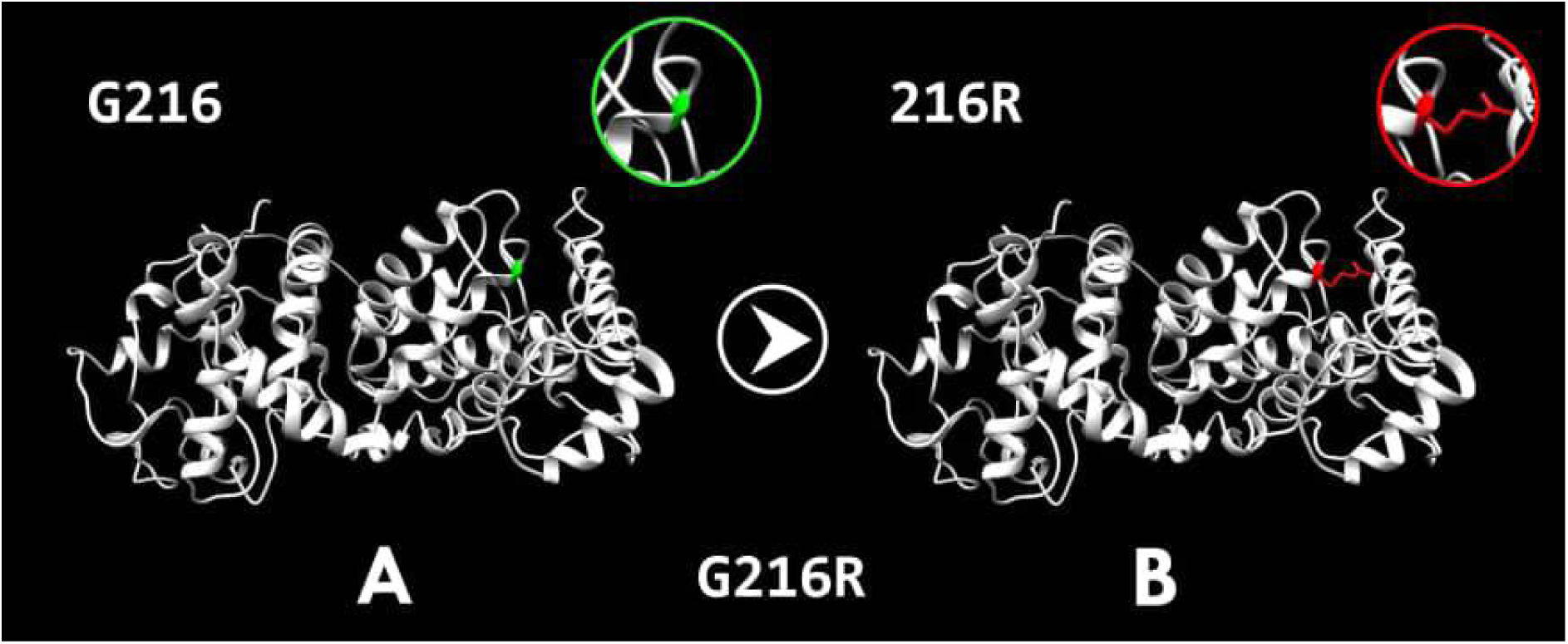
rs1579584286 (G216R). The amino acid Glycine in the wild protein (A) with green colour changed to Arginine in the mutant protein (B) with red colour at position 216.

The Y221C mutation replaces a large tyrosine with a smaller cysteine, potentially eliminating critical hydrophobic and aromatic interactions needed for structural stability. The W107C and L119V mutations also introduce smaller residues, which are predicted to create cavities and weaken essential contacts within the RHD structure (Figures 4, 5, and 6).

**Figure 4:**
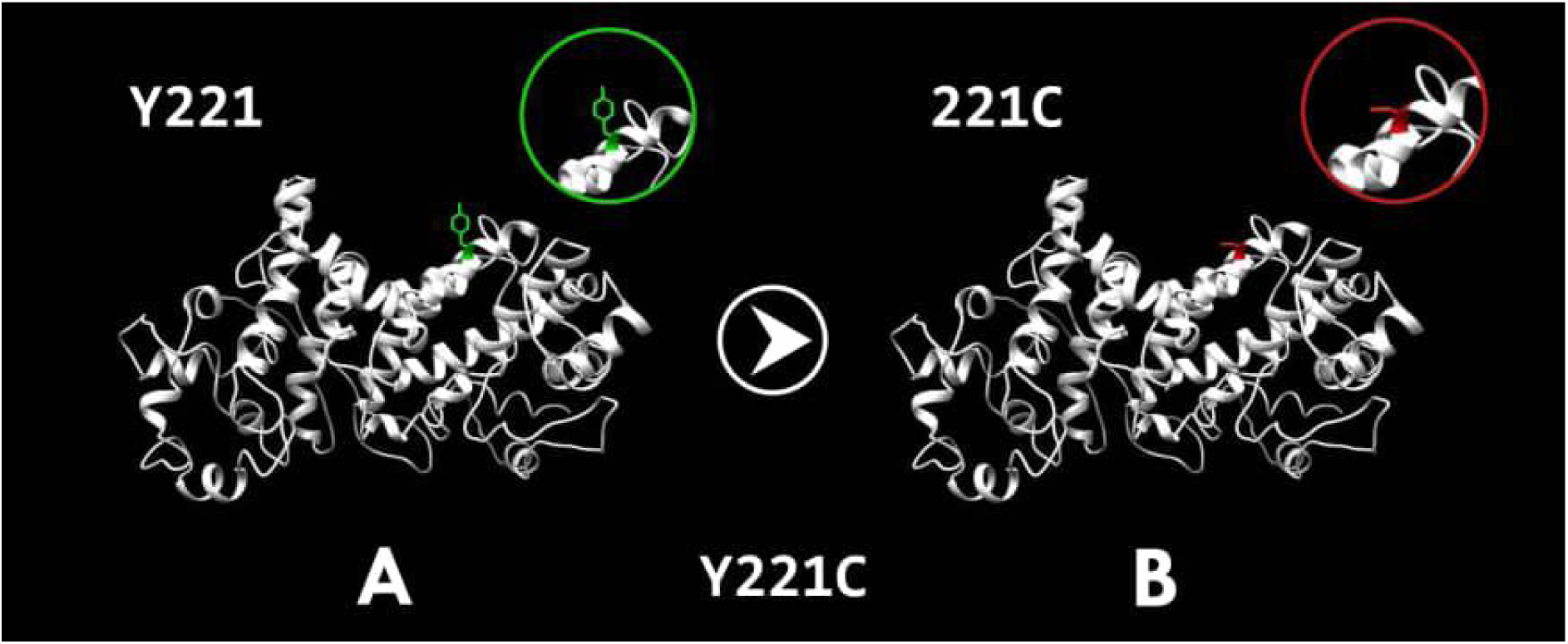
rs770744307 (Y221C). The amino acid Tyrosine in the wild protein (A) with green colour changed to cysteine in the mutant protein (B) with red colour at position 221.

**Figure 5:**
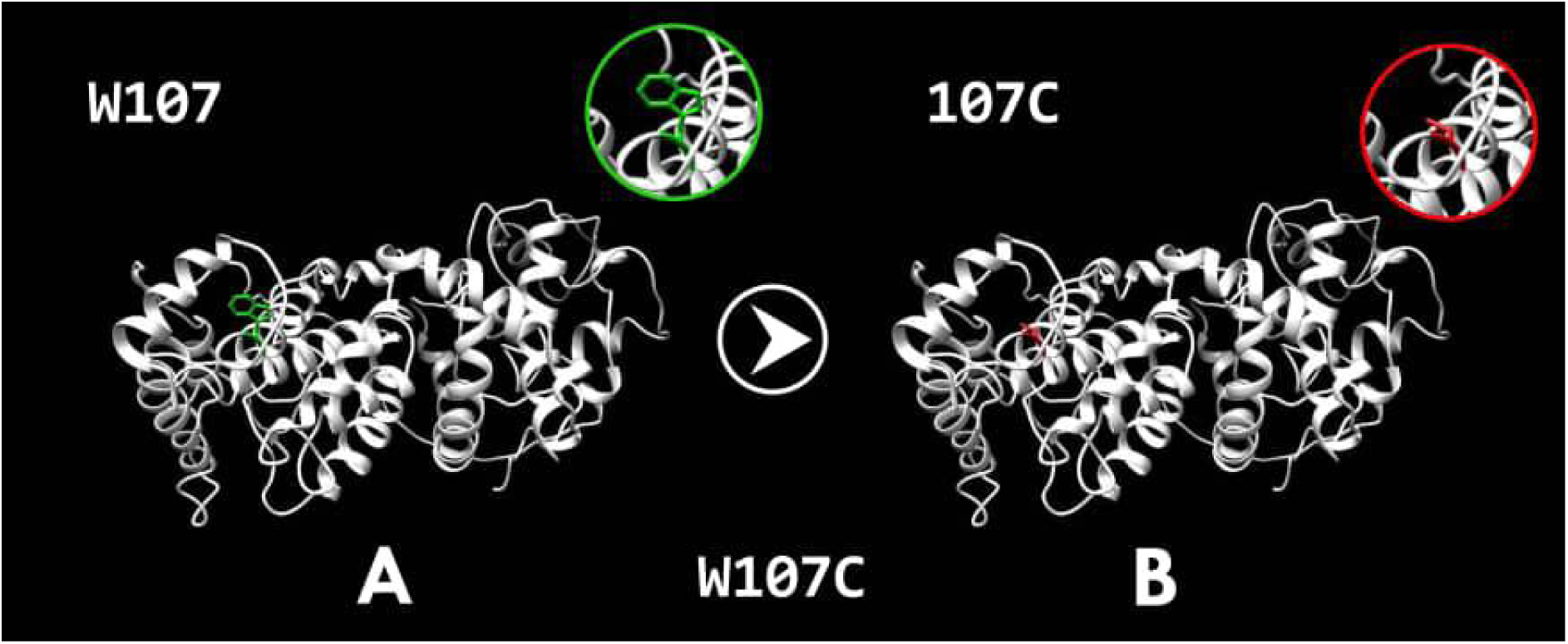
rs200065908 (W107C). The amino acid Tryptophan in the wild protein (A) with green colour changed to Cysteine in the mutant protein (B) with red colour at position 107.

**Figure 6:**
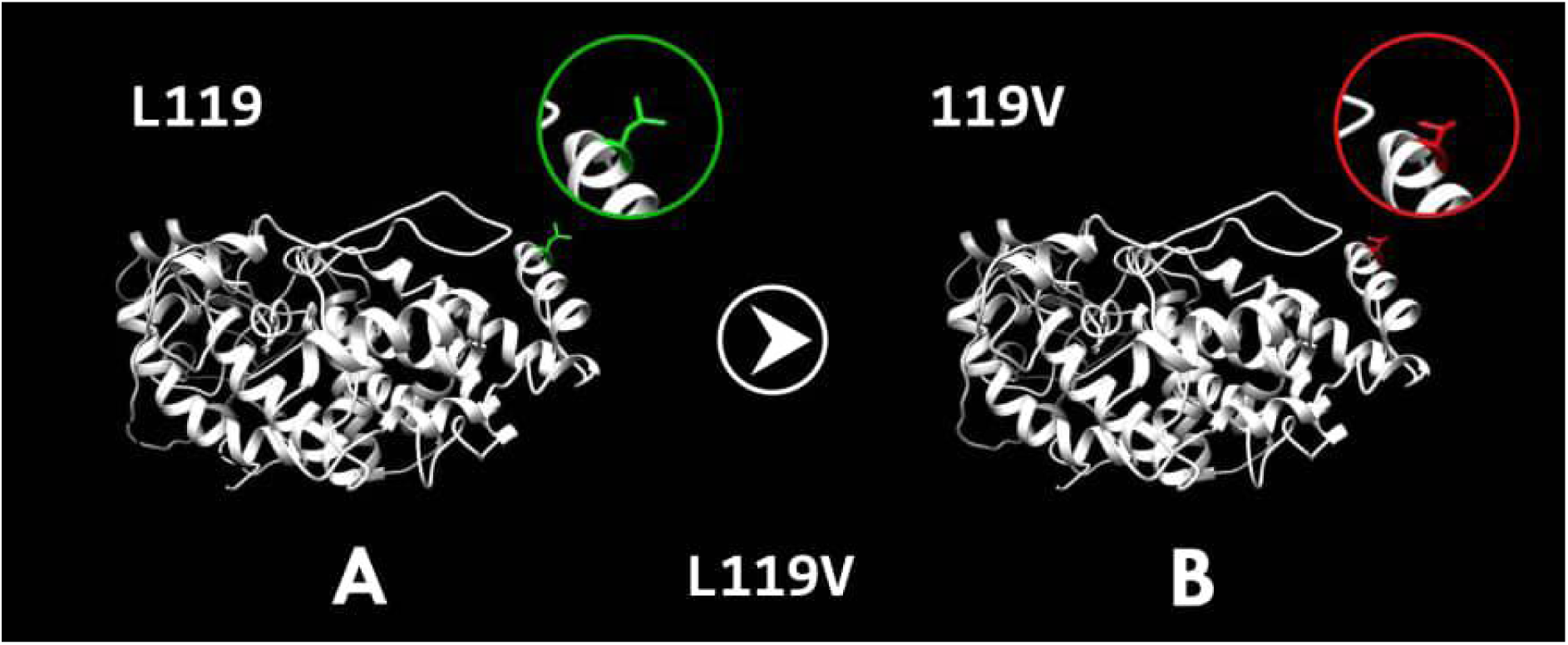
rs1467620688 (L119V). The amino acid Leucine in the wild protein (A) with green colour changed to Valine in the mutant protein (B) with red colour at position 119.

These structural perturbations provide a mechanistic basis for the predicted deleterious and destabilizing effects of these variants, suggesting they directly compromise the RHD’s core function.

### Protein Interaction Network and Functional Enrichment

#### Gene interaction(GeneMANIA)

To understand the functional context of RETREG1, we generated a protein-protein interaction network using GeneMania (Figure 7). The analysis revealed that RETREG1 has many vital function including cellular response to nitrogen levels, autophagosome, cellular response to starvation, organelle disassembly, response to starvation, mitochondrion disassembly, macroautophagy, vacuole organization, cellular response to nutrient levels, cellular response to extracellular stimulus, autophagosome organization, cellular response to external stimulus, response to nutrient levels, response to extracellular stimulus, ubiquitin-like protein ligase binding, and tubulin binding. The gene co-expressed with 12 genes, and physical interact with 26 genes. Physically, RETREG1 showed strong interactions with core autophagy proteins, most notably members of the GABARAP and MAP1LC3 (LC3) families, which are essential for autophagosome formation and maturation. This network strongly supports the role of RETREG1 in ER-phagy and suggests that deleterious mutations would disrupt this critical cellular pathway. (Figure, 7, and Supplementary table S1).

**Figure 7:**
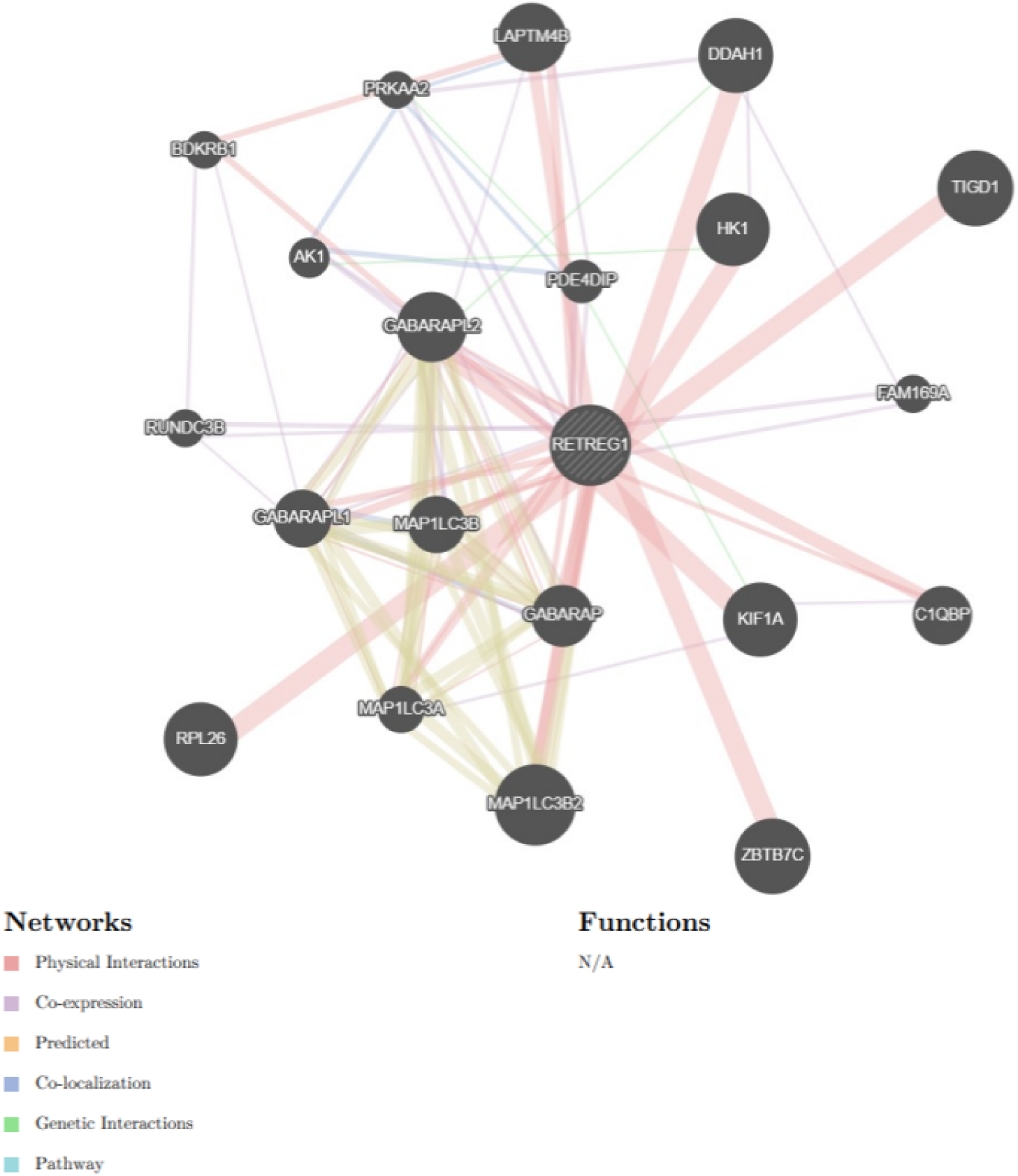
RETREG1 Gene interactions and network predicted by GeneMania. The meanings of the colours are shown at the bottom left

## Discussion

In the present study, we performed a comprehensive in silico analysis to identify and characterize potentially pathogenic non-synonymous single nucleotide polymorphisms (nsSNPs) in the RETREG1 gene associated with Hereditary Sensory and Autonomic Neuropathy type IIB (HSAN IIB). By systematically analyzing 345 reported nsSNPs using multiple complementary bioinformatics tools, we prioritized five high-confidence deleterious variants—W107C, L119V, G211R, G216R, and Y221C—that are predicted to significantly alter the structure, stability, and function of the RETREG1 protein.

The RETREG1 protein is crucial for maintaining endoplasmic reticulum (ER) health through its regulation of ER-phagy, a selective autophagic process that removes damaged or excess ER components (Berkane et al., 2023; Cinque et al., 2020; Kohno et al., 2019; Liao et al., 2019; Mo et al., 2020; Reggio et al., 2021; Wang et al., 2021). In addition, it also safeguards the ER homeostasis and preserves proper ER structure (Khaminets et al., 2015; Yang et al., 2021). Importantly, it also limits the accumulation of toxic byproducts of protein synthesis by mediating the autophagy-dependent lysosomal degradation of ER proteins (Chen et al., 2022).

RETREG1 has a two distinct domains, which are RHD domain and LIR domain (Gonzalez et al., 2023). RETREG1 RHD (residues of amino acids 80–240) (Popelka *et al.,* 2020). A common domain of selective autophagy receptors is the LIR motif (residues 453–458, C-terminus), which interacts to autophagy modifier proteins LC3/GABARAP family (Mo *et al.,* 2020). RETREG1 generates the ER membrane curvature using its RHD domain (Jiang et al., 2020).

All five high-risk mutations are located within the protein’s Reticulon Homology Domain (RHD), which is essential for shaping the ER membrane and facilitating the protein’s self-assembly (oligomerization), underscoring its non-redundant role in RETREG1 function. Therefore, we expect that all functions will be affected.

Research over the years has revealed the involvement of RETREG1 in numerous cancers, such as hepatocellular carcinoma (HCC) (Soner, 2013; Wang et al., 2021; Xie et al., 2023; YILMAZ & ÖZTÜRK, 2011), colorectal cancer (CRC) (Ajithkumar et al., 2024), breast cancer (Chipurupalli et al., 2022), and esophageal squamous cell carcinoma (ESCC) (Mo et al., 2020). Also RETREG1 functions as a tumor suppressor in cancer (Lee et al., 2020), preventing cancer from proliferating both *in vivo* and *in vitro* (Islam et al., 2018).

Although the precise molecular mechanisms underlying its tumor-suppressive role remain incompletely understood, alterations in RETREG1 expression or function appear to contribute to tumorigenesis.

In the present study, all five high-risk nsSNPs (W107C, L119V, G211R, G216R, and Y221C) were located within highly conserved regions of the RETREG1 protein. Given that evolutionary conservation often reflects functional importance, these mutations may impair RETREG1 activity and potentially compromise its tumor-suppressive function. However, the possible contribution of these variants to cancer development remains speculative and warrants further experimental and clinical investigation.

Since it may be difficult to interpret novel variants with clinical significance, numerous bioinformatics techniques have been developed to predict the biological effects of these polymorphisms. Using these techniques, the current study identified several unique SNPs in the RETREG1 protein and predicted the biological effects of these variants. Nevertheless, it is not possible to fully understand the functional importance of allelic variations using a single bioinformatics approach. Therefore, several complementary bioinformatics methods were employed in the present investigation. In silico analyses were used to identify the most likely functional variants in the RETREG1 gene, many of which remain uncharacterized.

The W107C variant was found in a region with moderate evolutionary conservation, and is predicted to be an exposed residue according to the NACSES algorithm, as indicated by ConSurf analysis.This observation supports findings reported in a previous study by Taşdelen et al., (2022) which identified a novel RETREG1 (FAM134B) allele associated with HSAN IIB and renal disease in a Turkish family. In that study, a unique homozygous ultrarare RETREG1 variant, NM_001034850.2:c.321G>A; p.Trp107Ter, was identified by exome sequencing. Electrophysiological examination of the proband revealed axonal sensorimotor neuropathy, predominantly affecting the lower limbs. Based on these findings, we also expect that the W107C variant may contribute to HSAN IIB pathology, as the wild-type residue at this position is highly conserved, with only a limited number of alternative residues observed across species.

The rs200065908 (W107C) mutation involves substitution of the wild-type tryptophan residue with cysteine, which is smaller in size. This size reduction may affect interactions with the lipid membrane and could lead to loss of critical contacts, as suggested by Project HOPE analysis. Furthermore, homologous sequences do not contain cysteine or residues with similar physicochemical properties at this position. According to conservation scores, this mutation is likely to be deleterious. The mutant residue is located near a highly conserved region, further supporting its potential functional importance. Additionally, MutPred analysis predicted that the W107C mutation may alter signal peptide–related properties.

Hereditary sensory and autonomic neuropathy is caused by RETREG1 dysfunction (Jiang et al., 2020). The RETREG1-G216R variant, on the other hand, appears to be a gain-of-function mutant, as it exhibits hyperactivity in oligomerization, ER fragmentation, and ER-phagy, ultimately leading to the death of sensory neurons. Remarkably. RETREG1-G216R also demonstrated significantly increased activity in liposome fragmentation, which is partially neutralized by the S149A, S151A, and S153A mutations. Taken together, RETREG1-G216R exhibits enhanced oligomerization activity, resulting in abnormal ER scission and increased ER-phagy (Jiang et al., 2020).

Mutations at position G216 cause an abnormal of RETREG1 oligomerization, which ultimately leads to HSAN II. Furthermore, in primary culture of dorsal root ganglion DRG sensory neurons, overexpression of RETREG1-G216R causes progressive cell death (Chen et al., 2022). This result (G216R) This finding for the G216R mutation is consistent with reports in the literature and further supports the hypothesis that this SNP may be a causative factor in HSAN IIB.

Project HOPE analysis predicted that the five highly deleterious nsSNPs identified in this study would negatively affect RETREG1 protein structure. Specifically, for rs1579584286 (G216R), substitution of glycine with arginine at position 216 introduces a residue that is larger than the wild-type glycine (Figure 4), which may disrupt interactions with the lipid membrane and result in steric clashes. In addition, the wild-type residue is neutral, whereas the mutant residue carries a positive charge. The wild-type residue is also more hydrophobic than the mutant residue; such differences in hydrophobicity may alter interactions with membrane lipids.

This mutation is located within a region annotated in UniProt as the reticulon homology domain (RHD), a functionally critical domain of RETREG1. Alterations in amino acid properties within this region may therefore disrupt its structure and function. Glycine, the wild-type residue at this position, is the most flexible amino acid, and this flexibility is likely essential for proper protein function. Substitution of this glycine residue may abolish this flexibility and impair protein activity. Moreover, mutation of a fully conserved residue is generally damaging to protein function. The mutant and wild-type residues differ substantially in their physicochemical properties, and based on conservation analysis, this mutation is therefore predicted to be deleterious. Additionally, the mutant residue is located near a highly conserved region, further supporting its functional importance.

As mentioned previously, to the best of our knowledge and based on available data, four of the five identified nsSNPs are novel variants. Consequently, interpretation of the remaining results relied primarily on Project HOPE predictions to elucidate the potential structural and functional consequences of these variants.

rs1467620688 (L119V) involves the substitution of leucine with valine at position 119. The mutant residue is smaller in size than the wild-type residue (Figure 6); therefore, interactions with the lipid membrane may be affected due to this size difference. This mutation is located within the RHD region. Differences in amino acid properties may disrupt this region and impair its function. The wild-type residue at this position is highly conserved; however, a limited number of other residue types have also been observed. The mutant residue was not among the residue types observed at this position in homologous proteins. Nevertheless, residues with some physicochemical properties similar to valine were identified, suggesting that in rare cases this mutation may occur without severely damaging protein function.

rs1738748948 (G211R) results from the substitution of glycine with arginine at position 211. The mutant residue is larger than the wild-type residue (Figure 5); therefore, interactions with the lipid membrane are likely to be affected, potentially leading to steric clashes. The wild-type residue is electrically neutral, whereas the mutant residue carries a positive charge, which may cause repulsion of ligands or neighboring residues with similar charges. In addition, the wild-type residue is more hydrophobic than the mutant residue, and differences in hydrophobicity may disrupt interactions with membrane lipids.

This residue is located within a region annotated in the UniProt database as a transmembrane domain, and only this residue type has been observed at this position. Mutation of a 100% conserved residue is generally damaging to protein function. The wild-type and mutant residues differ substantially in their physicochemical properties, and based on conservation analysis, this mutation is therefore predicted to be deleterious. Glycine, the wild-type residue, is the most flexible amino acid, and this flexibility is often essential for protein function. Substitution of glycine may abolish this flexibility, as glycine alone can adopt unusual torsion angles. Mutation to another residue may force the local backbone into an unfavorable conformation and disrupt local structure.

rs770744307 (Y221C) involves the substitution of tyrosine with cysteine at position 221. The mutant residue is smaller than the wild-type residue (Figure 3); thus, interactions with the lipid membrane may be affected, potentially leading to loss of interactions. In addition, the mutant residue is more hydrophobic than the wild-type residue. Differences in hydrophobicity may alter hydrophobic interactions with membrane lipids and lead to disruption of hydrogen bonds and/or improper protein folding.

This residue is located within a region annotated in the UniProt database as a transmembrane domain and lies within a stretch of residues known as the reticulon homology domain (RHD), which is identified in UniProt as a special region. Differences in amino acid properties may therefore disrupt this region and impair its function. At this position, only one residue type has been observed across homologous sequences. Mutation of a fully conserved residue typically has deleterious effects on protein function. The mutant and wild-type residues are not highly similar, and based on conservation analysis, this mutation is predicted to be damaging. Furthermore, the mutant residue is located near a highly conserved region, supporting its functional significance

GeneMANIA predicted that RETREG1 is involved in several biological processes, including cellular responses to nitrogen and nutrient levels, autophagosome formation and organization, cellular response to starvation, organelle and mitochondrion disassembly, macroautophagy, vacuole organization, cellular response to extracellular stimuli, ubiquitin-like protein ligase binding, and tubulin binding.

The present study is computational in nature. Although the concordance among multiple prediction tools provides strong supporting evidence, the pathogenicity of these novel variants must ultimately be validated through functional assays in cellular and animal models, as well as through genetic screening in HSAN IIB patient cohorts.

## Conclusion

This study employed an integrated in silico approach to evaluate the structural and functional impact of non-synonymous SNPs in the RETREG1 gene associated with HSAN IIB. Five high-risk variants (W107C, L119V, G211R, G216R, and Y221C) were identified, all of which localize within the functionally critical reticulon homology domain (RHD). Computational analyses indicate that these mutations may disrupt protein stability, membrane interactions, and ER-phagy-related functions. The detection of the known pathogenic G216R variant validates the analytical strategy, while the remaining four variants represent potentially novel deleterious mutations.

Overall, these findings expand the mutational spectrum of RETREG1 and provide mechanistic insight into HSAN IIB pathogenesis. Although based on computational predictions, this work establishes a framework for future experimental validation and genetic screening studies to confirm the clinical relevance of these variants.

---

**Supplementary table S1:**
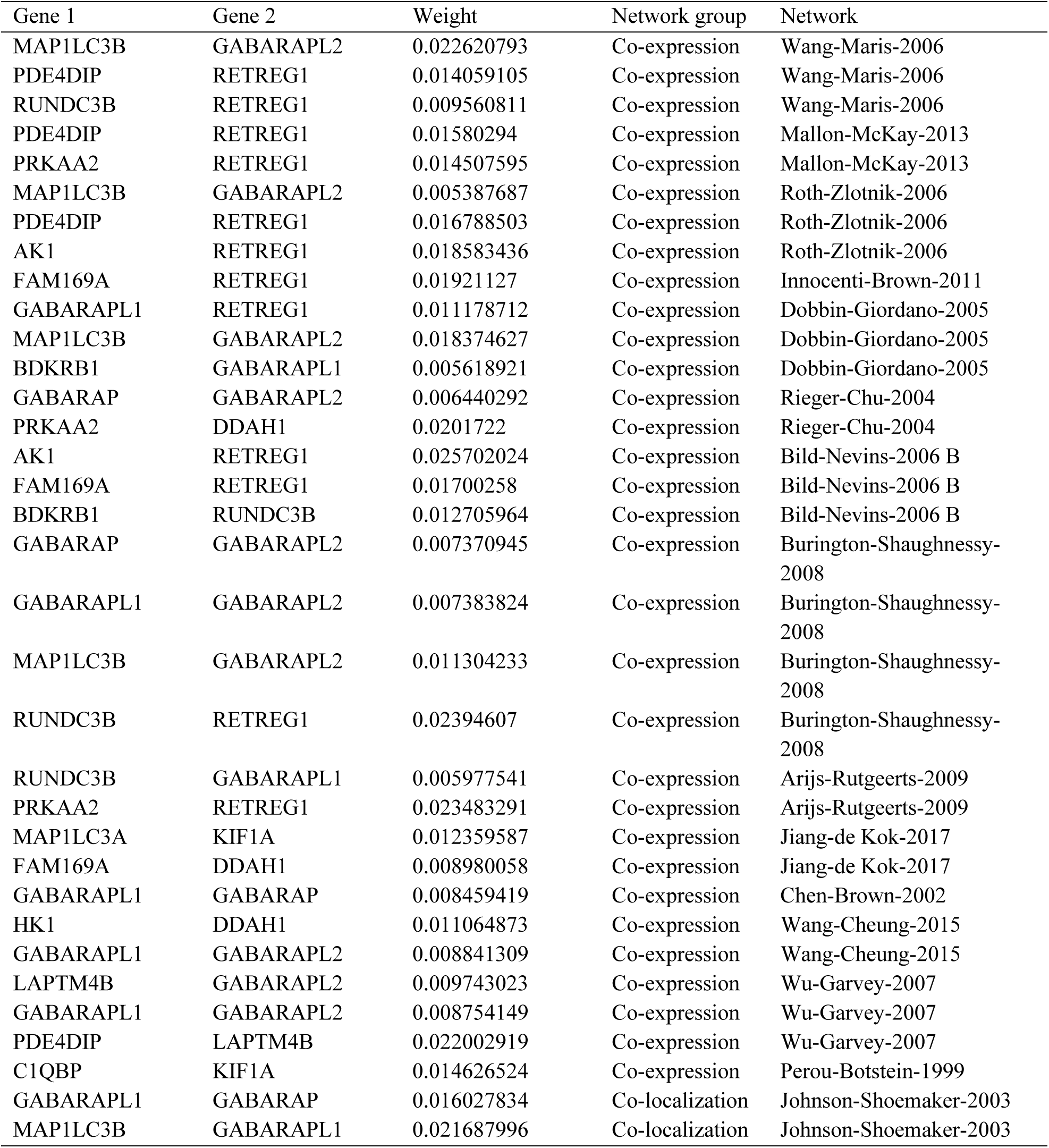
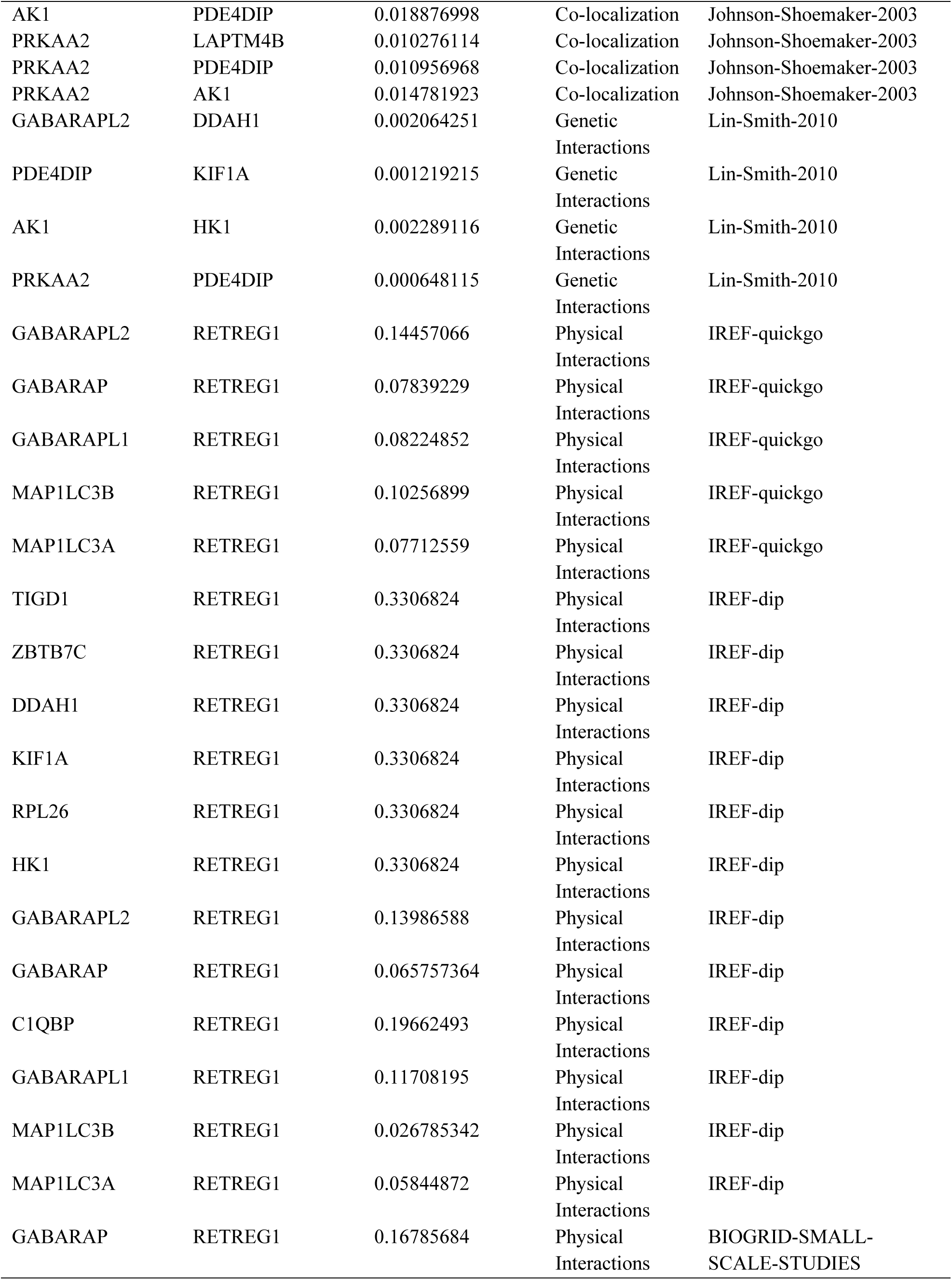
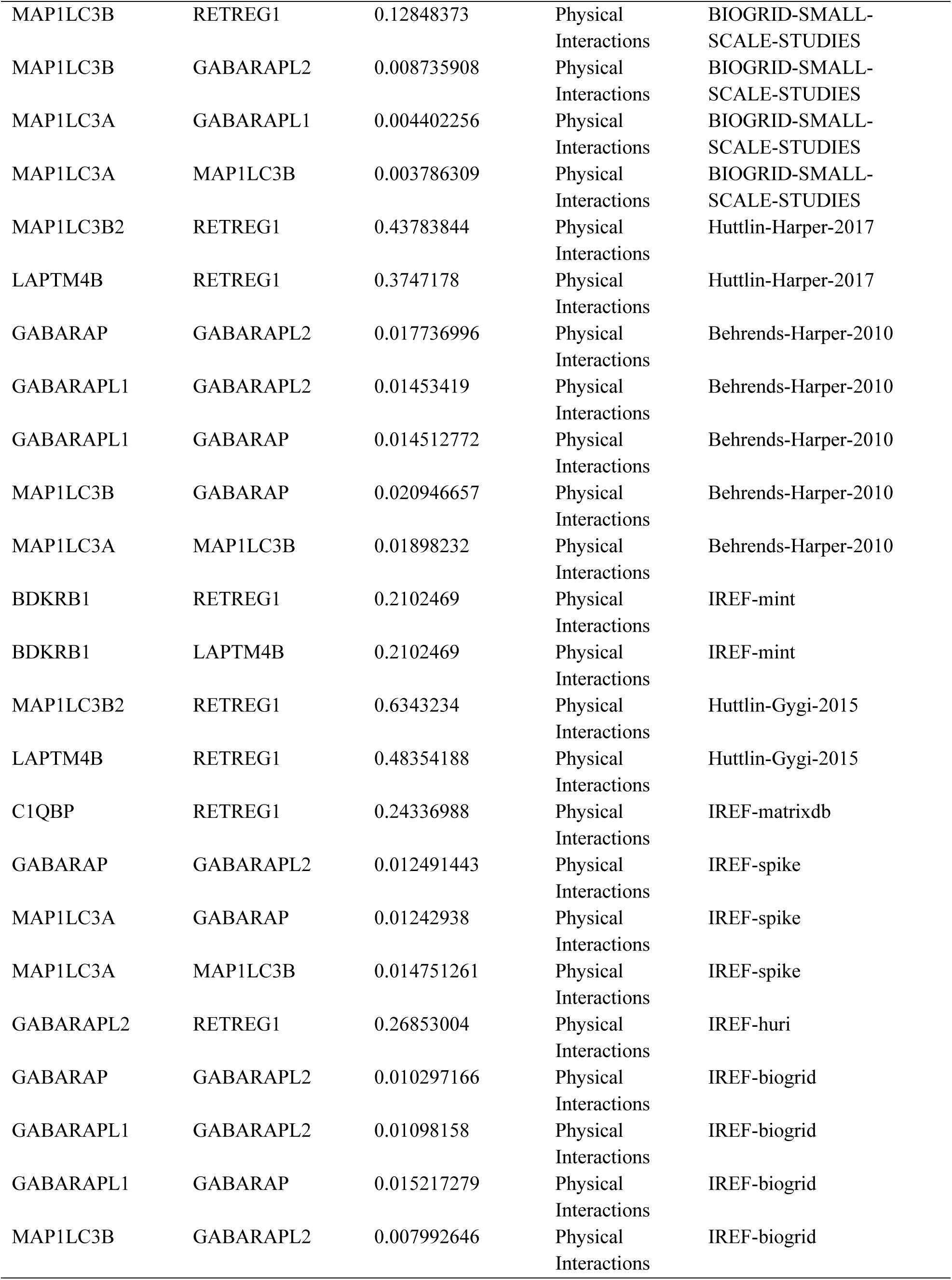
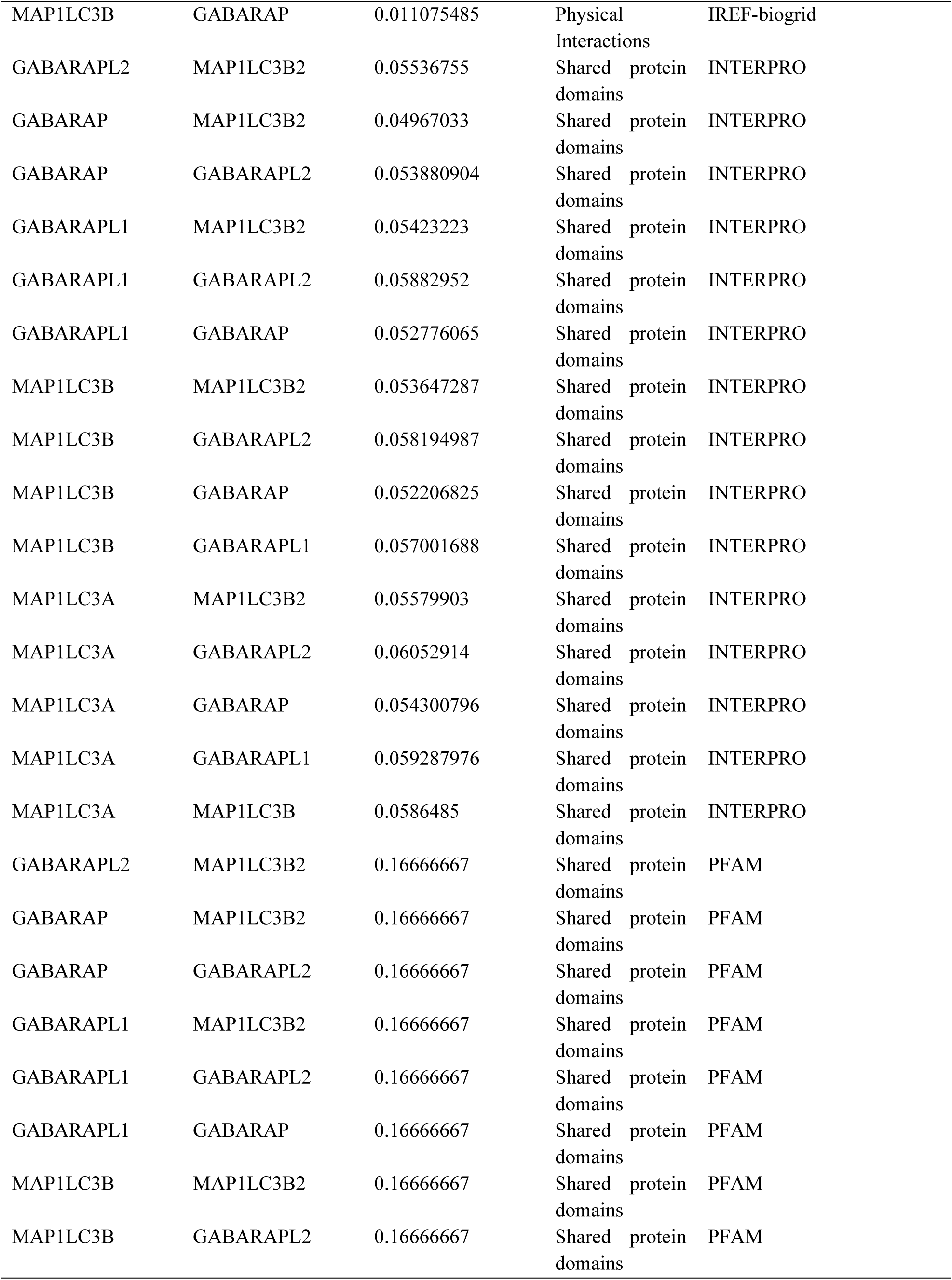
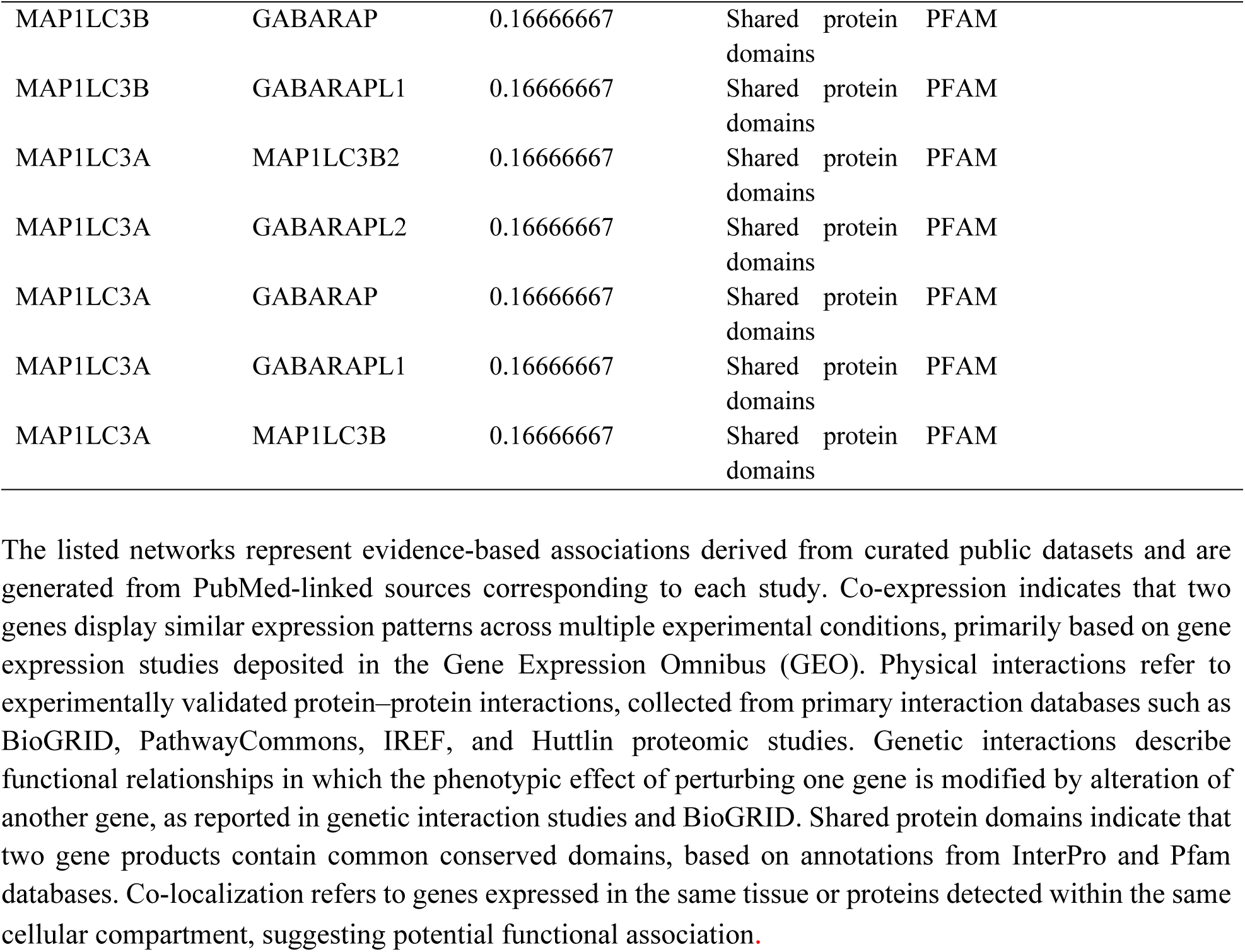
Gene network analysis showing genes that are co-expressed with, share protein domains with, co-localize with, or physically and genetically interact with RETREG1.

